# HIV-1 accessory protein Vpr interacts with REAF/RPRD2 to mitigate its antiviral activity

**DOI:** 10.1101/408161

**Authors:** Joseph M Gibbons, Kelly M Marno, Rebecca Pike, Wing-yiu Jason Lee, Christopher E Jones, Babatunji W Ogunkolade, Claire Pardieu, Alexander Bryan, Rebecca Menhua Fu, Gary Warnes, Paul A Rowley, Richard D Sloan, Áine McKnight

## Abstract

The Human Immunodeficiency Virus type 1 (HIV-1) accessory protein Vpr enhances viral replication in both macrophages and in cycling T cells to a lesser extent. Virion packaged Vpr is released in target cells shortly after entry, suggesting its requirement in the early phase of infection. Previously, we described REAF (RNA-associated Early-stage Antiviral Factor, RPRD2), a constitutively expressed protein that potently restricts HIV replication at or during reverse transcription. Here, we show that a virus without intact *vpr* is more highly restricted by REAF and, using delivery by VLPs, that Vpr alone is sufficient for REAF degradation in primary macrophages. REAF is more highly expressed in macrophages than in cycling T cells and we detect, by co-immunoprecipitation assay, an interaction between Vpr protein and endogenous REAF. Vpr acts very quickly during the early phase of replication and induces the degradation of REAF within 30 minutes of viral entry. Using Vpr F34I and Q65R viral mutants, we show that nuclear localisation and interaction with cullin4A-DBB1 (DCAF1) E3 ubiquitin ligase is required for REAF degradation by Vpr. In response to infection, cells upregulate REAF levels. This response is curtailed in the presence of Vpr. These findings support the hypothesis that Vpr induces the degradation of a factor, REAF, which impedes HIV infection in macrophages.

**Importance:** For at least 30 years, it has been known that HIV-1 Vpr, a protein carried in the virion, is important for efficient infection of primary macrophages. Vpr is also a determinant of the pathogenic effects of HIV-1 *in vivo*. A number of cellular proteins that interact with Vpr have been identified. So far, it has not been possible to associate these proteins with altered viral replication in macrophages, or to explain why Vpr is carried in the virus particle. Here we show that Vpr mitigates the antiviral effects of REAF, a protein highly expressed in primary macrophages and one which inhibits virus replication early during reverse transcription. REAF is degraded by Vpr within 30 minutes of virus entry, in a manner dependent on the nuclear localization of Vpr and its interaction with the cell’s protein degradation machinery.

## Introduction

Human Immunodeficiency Virus type 1 (HIV-1) infects CD4^+^ T cells and macrophages *in vivo*. HIV-1 has four non-structural accessory genes *nef*, *vif*, *vpu* and *vpr*. *Nef, vif and vpu* diminish host innate immunity. A function for Vpr has been elusive, but it is required for efficient replication in macrophages and for pathogenesis *in vivo* (1, 2). A widely acknowledged but poorly understood Vpr-mediated phenotype is that it induces cell cycle arrest at the G2/M phase using the cullin4A-DBB1 (DCAF1) E3 ubiquitin ligase and the recruitment of an unknown substrate for proteasomal degradation. A large number of Vpr substrates have been reported (3–11). Yan *et al.* (2019) show that helicase-like transcription factor (HLTF) weakly restricts replication of HIV-1 in T cells. HLTF was shown previously to be down modulated by Vpr (12, 13). Furthermore, Greenwood *et al.* 2019 report that Vpr promotes large scale remodelling of approximately 2000 cellular proteins, including those that bind nucleic acids and ones involved with the cell cycle (14).

Substantial quantities of Vpr are incorporated into viral particles and released from the major capsid protein (CA) after entry into the cell (15, 16). The timing of Vpr release coincides with the initiation of reverse transcription, a process that transcribes the RNA genome into DNA for subsequent integration into the host cell DNA (17). The early release of Vpr from the CA implies it has an early function prior to integration events. When considering the role of Vpr in cell tropism and pathogenesis, the investigation of proteins that have a direct effect on viral replication is a priority.

Here we focus on <u>R</u>NA-associated <u>E</u>arly-stage <u>A</u>ntiviral <u>F</u>actor (REAF, also known as <u>R</u>egulation of nuclear <u>p</u>re-m<u>R</u>NA <u>d</u>omain-containing protein <u>2</u>/RPRD2), originally described as a restriction to HIV replication and called lentiviral restriction 2 (Lv2) (18, 19). Lv2 was first shown to be a restriction to the replication of HIV-2 and subsequently it was shown to inhibit the replication of HIV-1 and SIV during reverse transcription (20). Lv2/REAF restriction is cell type dependent (19, 21–24), active in certain cell types including HeLa-CD4 and primary macrophages (18, 19). Susceptibility of the virus to Lv2 is determined by both the viral envelope (Env) and capsid (CA) (23, 24). REAF was identified in a whole genome siRNA screen for the identification of HIV-1 restriction factors. Like Lv2, REAF limits the populations of target cells completion of proviral DNA synthesis and integration of the viral genome (18). Subsequently, REAF was demonstrated to form a major component of Lv2 (19).

Here, we show that within 30 minutes of cellular entry, only HIV-1 virus that contains Vpr can induce the degradation of REAF and rescue efficient viral replication in primary macrophages. Using Vpr mutant viruses, we demonstrate that the nuclear localisation of Vpr, and its ability to interact with cullin4A-DBB1 (DCAF1) E3 ubiquitin ligase, is a requirement for REAF degradation. Down modulation of REAF by Vpr in the early phase of infection is transient and re-expression to basal levels is achieved by approximately one hour. After infection with HIV-1, or treatment with polyriboinosinic:polyribocytidylic acid (poly(I:C)) or Lipopolysaccharide (LPS), cells respond by increasing REAF levels. In the case of viral infection, this response is curtailed in the presence of Vpr. Therefore, our results support the hypothesis that Vpr induces the degradation of a cellular protein, REAF, a protein which impedes HIV-1 infection in macrophages during reverse transcription.

## Results

### HIV-1 Vpr interacts with REAF and overcomes restriction

REAF restricts HIV-1 replication in HeLa-CD4 (18, 20). We sought to determine if a viral accessory gene could overcome REAF and so we tested the infectivity of HIV-1 89.6^WT^ and mutants deleted for *vpr* (89.6^Δ*vpr*^), *vif* (89.6^Δ*vif*^) or *vpu* (89.6^Δ*vpu*^) in these cells. Preventing REAF expression using short-hairpin RNA (HeLa-CD4 shRNA-REAF, Figure 1A) reveals its potent antiviral effect. Despite a standard input for each virus (50 FFU/ml, as measured on HeLa-CD4), there is significantly greater rescue of HIV-1 89.6^Δ*vpr*^ (>3 fold, p < 0.0001) compared to HIV-1 89.6^WT^ (Figure 1B). The prevention of REAF expression using shRNA alleviates the need for Vpr. Conversely, there was no significantly greater rescue for either HIV-1 89.6^Δ*vif*^ or HIV-1 89.6^Δ*vpu*^ compared to HIV-1 89.6^WT^ (data not shown). Thus, *vpr* potently overcomes the restriction imposed by REAF.

**Figure 1:**
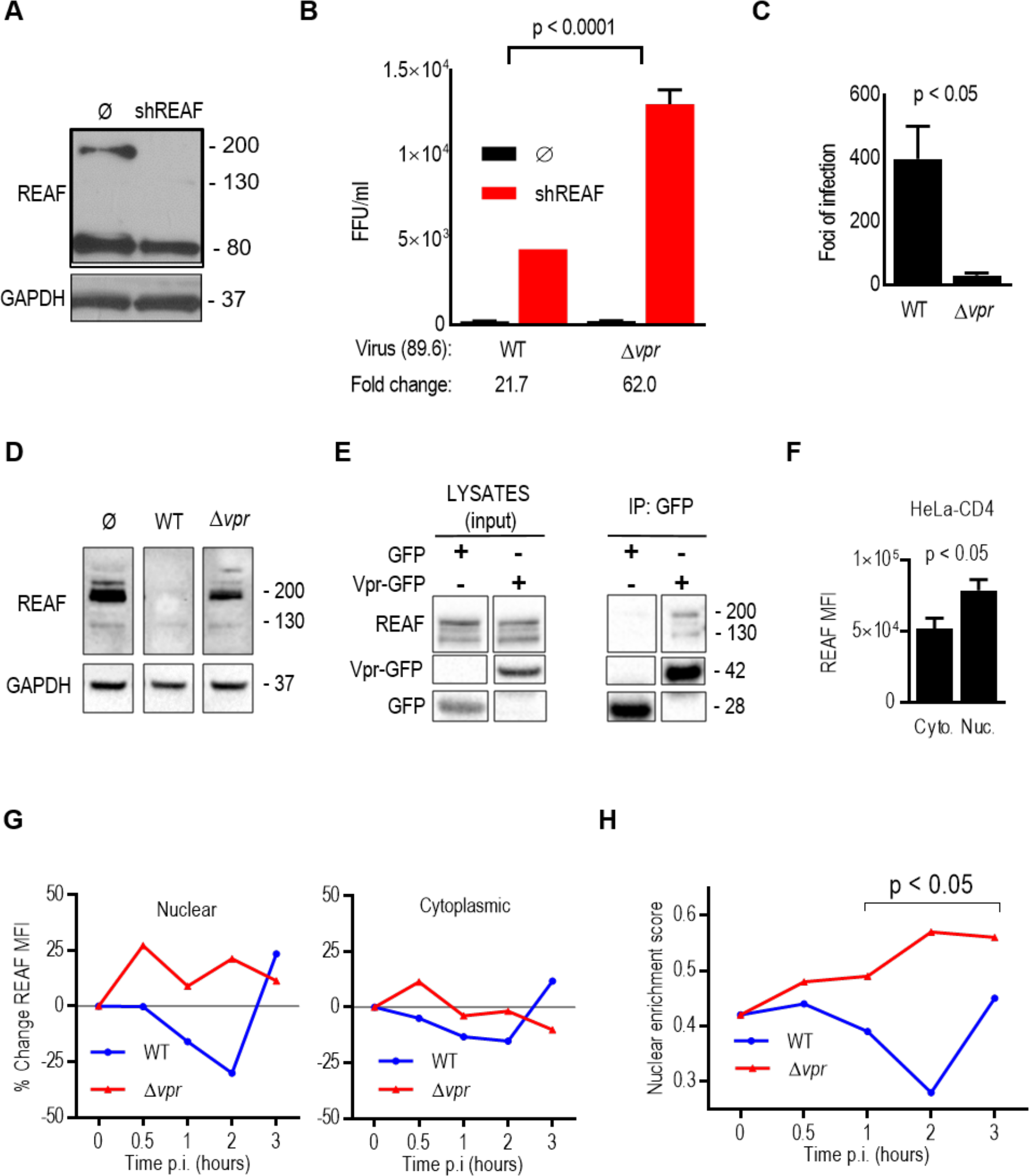
HIV-1 Vpr interacts with REAF and overcomes restriction. **(A)** REAF protein in HeLa-CD4 (Ø) and HeLa-CD4 shRNA-REAF (shREAF). GAPDH is a loading control. **(B)** Infectivity (FFU/ml) of HIV-1 89.6^WT^ and HIV-1 89.6^Δ*vpr*^ in HeLa-CD4 (Ø) and HeLa-CD4 shRNA-REAF (shREAF). Viral inputs were equivalent at approximately 50 FFU/ml measured on HeLa-CD4. Error bars in indicate the standard deviations of means derived from a range of duplicate titrations. Fold changes in FFU are indicated. **(C)** Resulting foci of infection from equal p24 inputs (1ng) of HIV-1 89.6^WT^ or HIV-1 89.6^Δ*vpr*^ in HeLa-CD4. Error bars in indicate the standard deviations of means derived from a range of duplicate titrations. **(D)** REAF protein in Hela-CD4 24 hours post challenge with HIV-1 89.6^WT^ or HIV-1 89.6^Δ*vpr*^. GAPDH is a loading control. **(E)** HEK-293T cells were transfected with Vpr-GFP expression plasmid or GFP control vector, expression was analysed by Western blotting (left) and protein was immunoprecipitated (IP) with anti-GFP beads. Co-immunoprecipitated REAF was detected in the Vpr-GFP precipitation (right). **(F)** Nuclear and cytoplasmic REAF mean fluorescence intensity (MFI) in HeLa-CD4 measured by imaging flow cytometry. Error bars represent standard deviations of means of replicates. **(G)** Percentage (%) change in REAF MFI from time ‘0’ in the nucleus (left) and cytoplasm (right) of Hela-CD4 over time after challenge with HIV-1 89.6^WT^ or HIV-1 89.6^Δ*vpr*^. Results are representative of three independent experiments. **(H)** Nuclear enrichment score of HeLa-CD4 over time post challenge with HIV-1 89.6^WT^ or HIV-1 89.6^Δvpr^. A lower nuclear enrichment score indicates a lower proportion of overall REAF is located in the nucleus as calculated by IDEAS software. Results are representative of three independent experiments. Where quantitative comparisons are made, blots are derived from the same blot or blots processed together.

We are unaware of previous reports that Vpr overcomes known or unknown HIV-1 restrictions in HeLa-CD4. Therefore, we confirmed that the mutant HIV-1 89.6^Δ*vpr*^ is restricted in HeLa-CD4 compared to HIV-1 89.6^WT^. Figure 1C shows that despite equal viral inputs (measured by ELISA of viral protein p24), significantly fewer foci of infection (FFU) result from challenge with HIV-1 89.6^Δ*vpr*^ compared to HIV-1 89.6^WT^. Further support of a role for Vpr in overcoming REAF is evidenced in Figure 1D. When HeLa-CD4 are challenged with a HIV-1 89.6^WT^ (which has an intact *vpr*), REAF protein is down modulated. The observed down modulation is dependent on the presence of Vpr as HIV-1 89.6^Δ*vpr*^ is incapable of degrading REAF. Moreover, Figure 1E shows that Vpr and REAF interact with each other, either directly or indirectly as part of a complex, as they are co-immunoprecipitated.

Vpr is released from the capsid and enters the nucleus shortly after infection (11). Imaging flow cytometry combines traditional flow cytometry with microscopy, facilitating the evaluation of both the expression and subcellular localisation of proteins in large populations of cells (25, 26). Using imaging flow cytometry, we determined the relative subcellular localization of REAF in HeLa-CD4. This analysis reveals that REAF is more highly expressed in the nuclear region compared to the cytoplasmic region of cycling HeLa-CD4 (Figure 1F).

Previously, we reported that REAF affects the production of reverse transcripts early in infection and that at this critical time point, REAF is transiently down modulated in HeLa-CD4 (20). Also using image flow cytometry, we looked the subcellular distribution of REAF at the early time points following HIV-1 infection. Here, REAF protein was quantified by imaging flow cytometry in the cytoplasm and nucleus of HeLa-CD4 over the first 3 hours of infection with either HIV-1 89.6^WT^ or HIV-1 89.6^Δ*vpr*^. Following challenge with HIV-1 89.6^Δ*vpr*^, REAF levels increase within 0.5 hours in both the nucleus (~25%, Figure 1G, left) and cytoplasm (~10%, Figure 1G, right). Nuclear levels remain high for 3 hours. Conversely, in the presence of Vpr (HIV-1 89.6^WT^) this increase in REAF is curtailed, instead there is a steady decline from 0.5-2 hours. The decline is most apparent in the nucleus with ~20% reduction by 1 hour and ~30% at 2 hours. By 3 hours, levels of REAF protein recover. The virus carries limited quantities of Vpr (17), which potentially explains why there is a pause in REAF down modulation. Lower levels of REAF were also observed in the cytoplasm over time after infection with HIV-1 89.6^WT^, but to a much lesser extent (Figure 1G, right). The nuclear enrichment score (NES) is a comparison of the intensity of REAF fluorescence inside the nucleus (defined using DAPI) to the total fluorescence intensity of REAF in the entire cell (defined using brightfield images). The lower the score, the less REAF in the nucleus relative to in the cell overall. Imaging flow cytometry software determined the NES over time after infection with HIV-1 89.6^WT^ or HIV-1 89.6^Δ*vpr*^ (Figure 1H). By 1-2 hours, a significant segregation emerges; in the presence of Vpr relative nuclear levels of REAF are suppressed. The greatest segregation occurs 2 hours post infection.

### Fluctuations in subcellular REAF expression after HIV-1 infection are Vpr dependent

Macrophages are a target for HIV infection *in vivo* (27). Vpr has been shown, to varying degrees, to be more beneficial for replication in these cells than in cycling T cells (28–32). For that reason, we investigated REAF effects in monocyte-derived macrophages (MDMs). Also using imaging flow cytometry, we determined that similar to HeLa-CD4, MDMs have significantly greater quantities of REAF in the nucleus compared to in the cytoplasm (Figure 2A). Nuclear levels of REAF were also compared in a number of primary cell types using imaging flow cytometry (Figure 2B). When compared with either monocytes or resting/activated T cells, both MDMs and dendritic cells (DCs) highly express nuclear REAF. In Figure 2C, MDMs were treated with virus-like particles (VLPs) containing Vpr and Western blotting confirmed that Vpr down modulates REAF in MDMs and that Vpr alone is sufficient to induce this down modulation.

**Figure 2:**
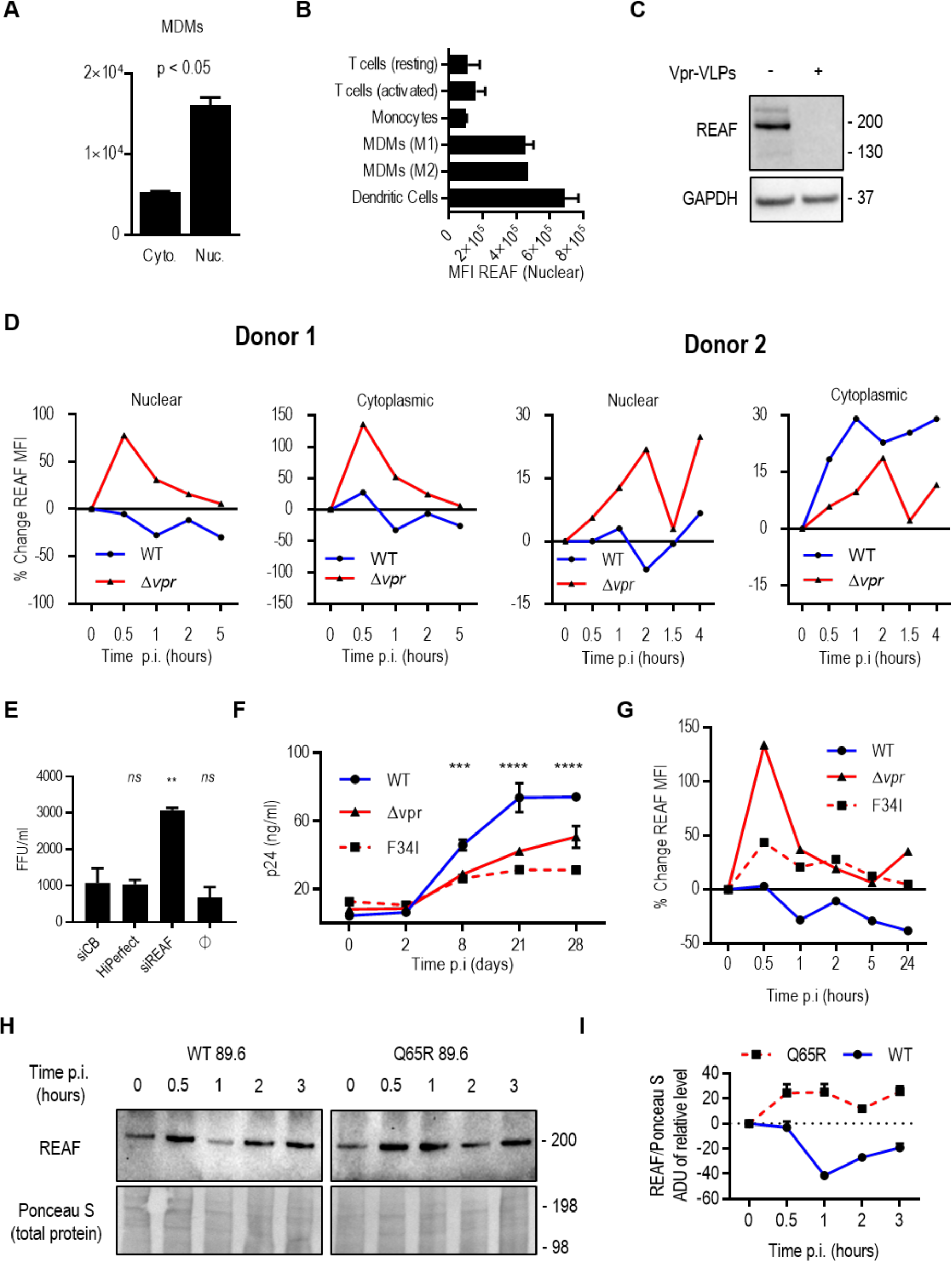
Fluctuations in subcellular REAF expression after HIV-1 infection are Vpr dependent. **(A)** Nuclear and cytoplasmic REAF mean fluorescence intensity (MFI) in MDMs measured by imaging flow cytometry. Error bars represent standard deviations of means of replicates. **(B)** Nuclear REAF MFI in indicated primary cell types measured by imaging flow cytometry. Error bars represent standard deviations of means of two blood donors. **(C)** REAF protein in MDMs treated with empty or Vpr-containing VLPs. GAPDH is a loading control. VLP input was equivalent at 100ng of p24. **(D)** Percentage (%) change in subcellular REAF MFI in MDMs from time ‘0’ measured by imaging flow cytometry after challenge with HIV-1 89.6^WT^ or HIV-1 89.6^Δ*vpr*^. Data from two donors are presented. **(E)** Infectivity (FFU/ml) of HIV-1 89.6^WT^ in MDMs transfected with siRNA-REAF. HiPerfect (transfection reagent) and siCB are negative controls. ** = P<0.01, *ns* = not significant, one-way ANOVA and *post hoc* Dunnett’s test. **(F)** Infectivity of HIV-1 89.6^WT^ compared with HIV-1 89.6^Δ*vpr*^ and HIV-1 89.6 ^F34I^ in MDMs. p24 antigen concentrations over 28 days post infection are indicated. Viral input was equivalent at 50ng of p24. Error bars represent standard deviations of means of duplicates (*** = P < 0.001; **** = P < 0.0001, two-way ANOVA; the same results were obtained for HIV-1 89.6^WT^ versus HIV-1 89.6^Δ*vpr*^ and HIV-1 89.6 ^F34I^). Data is representative of at least two independent experiments. **(G)** Percentage (%) change in total cellular REAF MFI from time ‘0’ in MDMs after challenge with HIV-1 89.6^WT^, HIV-1 89.6^Δ*vpr*^ or HIV-1 89.6^F34I^. Results are representative of three independent experiments. **(H)** REAF protein, measured by Western blotting, in MDMs challenged with HIV-1 89.6^WT^ **or** HIV-1 89.6^Q65R^ over time. Ponceau S staining of nitrocellulose membrane is a loading control. Associated densitometry is presented in **(I)** where error bars represent standard deviations of means where analysis was performed in triplicate.

We investigated the ability of Vpr to degrade REAF in MDMs early in infection. The subcellular fluctuations in REAF mean fluorescence intensity (MFI) were measured by imaging flow cytometry in large populations of target cells (>5000). In the presence of Vpr (HIV-1 89.6^WT^), nuclear REAF decreases within 2 hours of viral infection of macrophages from two donors (Figure 2D), similar to that observed in HeLa-CD4 (Figure 1G). In contrast, also in both donors, nuclear REAF rapidly increases from as early as 0.5 hours when the virus does not contain Vpr (HIV-1 89.6^Δ*vpr*^) (Figure 2D). For the cytoplasmic compartment, a similar picture emerges for REAF fluctuation. In both donors, when Vpr is absent, REAF levels increase rapidly within 0.5 hours of infection (Figure 2D). This cytoplasmic increase is curtailed in donor 1 when Vpr is present. In donor 2, the loss of nuclear REAF after HIV-1 89.6^WT^ infection is paralleled by an increase in cytoplasmic REAF. Similar kinetics of *total* REAF protein fluctuation were measured by Western blotting in the presence or absence of Vpr in MDMs from two further donors (data not shown).

We sought to determine if knockdown of REAF in primary macrophages results in an increase in susceptibility to HIV-1 infection. In Figure 2E, primary MDMs were treated with siRNA targeting REAF (siREAF) or a control protein (siCB). Cells lacking REAF were found to be significantly (p<0.0001) more susceptible to infection with HIV-1 89.6. We confirmed previous reports that HIV-1 replication in MDMs is more efficient in the presence of Vpr (27, 30). Figure 2F shows that HIV-1 89.6^Δ*vpr*^ has restricted replication in MDMs compared with the wild type virus expressing Vpr (HIV-1 89.6^WT^).

To investigate further the relationship between nuclear REAF and Vpr, we generated a virus with a substitution within Vpr (F34I). HIV-1 89.6^F34I^ is incapable of localising to the nuclear membrane or of interacting with the nuclear transport protein importin-α and nucleoporins (30). Like HIV-1 89.6^Δ*vpr*^, the mutant virus (HIV-1 89.6^F34I^) replicates less efficiently in MDMs (Figure 2F). Using imaging flow cytometry, the respective abilities of these three viruses (HIV-1 89.6^WT^, 89.6^Δ*vpr*^ and 89.6^F34I^) in down modulating total REAF protein was investigated in MDMs (Figure 2G). As expected, there is a loss of total REAF from 30 minutes after HIV-1 89.6^WT^ infection (Figure 2G) with a transient recovery at around 2 hours. The opposite occurs in the absence of Vpr (HIV-1 89.6^Δ*vpr*^), REAF levels *increase* after infection. The increase in REAF levels is most potent after 30 minutes of infection with HIV-1 89.6^Δ*vpr*^. HIV-1 89.6^F34I^, similar to HIV-1 89.6 ^Δ*vpr*^ can no longer deplete REAF in MDMs (Figure 2G).

Other targets of Vpr have been reported. It recruits SLX4-SLX1/MUS81-EME1 endonucleases to DCAF1, activating MUS81 degradation and triggering arrest in G2/M (33). It also degrades helicase-like transcription factor (HLTF), a protein recently shown to enhance infection of HIV-1 in T-cell lines (8, 12). To determine if the depletion of REAF requires the association of Vpr with DCAF1, we generated another mutant virus with a different substitution within *vpr*, Q65R. Previously, the Q65R mutation was shown to ablate the association between DCAF1 and Vpr and the ability of Vpr to induce arrest at G2/M (34–36). Figures 2H and I show that this mutant, compared to HIV-1 89.6^WT^, is unable to down modulate REAF. We cannot rule out inhibition of synthesis or increased nuclear export in addition to degradation as possibilities.

### Expression of REAF during cell cycle

A phenotype of HIV-1 Vpr is that it can induce cell cycle arrest at G2/M in cycling T cells (27, 37–39). The failure of both HIV-1 89.6^F34I^ and HIV-1 89.6^Q65R^ to efficiently induce G2/M arrest (30, 34, 40–42), and our observation that they cannot down modulate REAF (Figures 2G, H and I), prompted us to investigate REAF and the cell cycle.

First, we determined the expression levels of REAF at various phases of the cell cycle using imaging flow cytometry (Figure 3A). REAF protein levels are lowest in G0/1, increase through S phase, and peak in G2/M. Confocal microscopy of cycling cells concurred with the quantitative analysis in Figure 3A, overall REAF levels appeared greater in mitotic cells (Figure 3B). There is also an apparent exclusion of REAF from the nuclear region of the cell during mitosis (particularly during metaphase, anaphase and telophase). Quantitative analysis by imaging flow cytometry of cycling cells confirmed that the mitotic population had a lower nuclear enrichment score (0.13) compared to the non-mitotic cells (1.53), indicating a lower intensity of REAF in the nucleus compared to in the cell as a whole (Figure 3C). Representative images of subcellular REAF in mitotic and non-mitotic cells from imaging flow cytometry are presented in Figure 3D.

**Figure 3:**
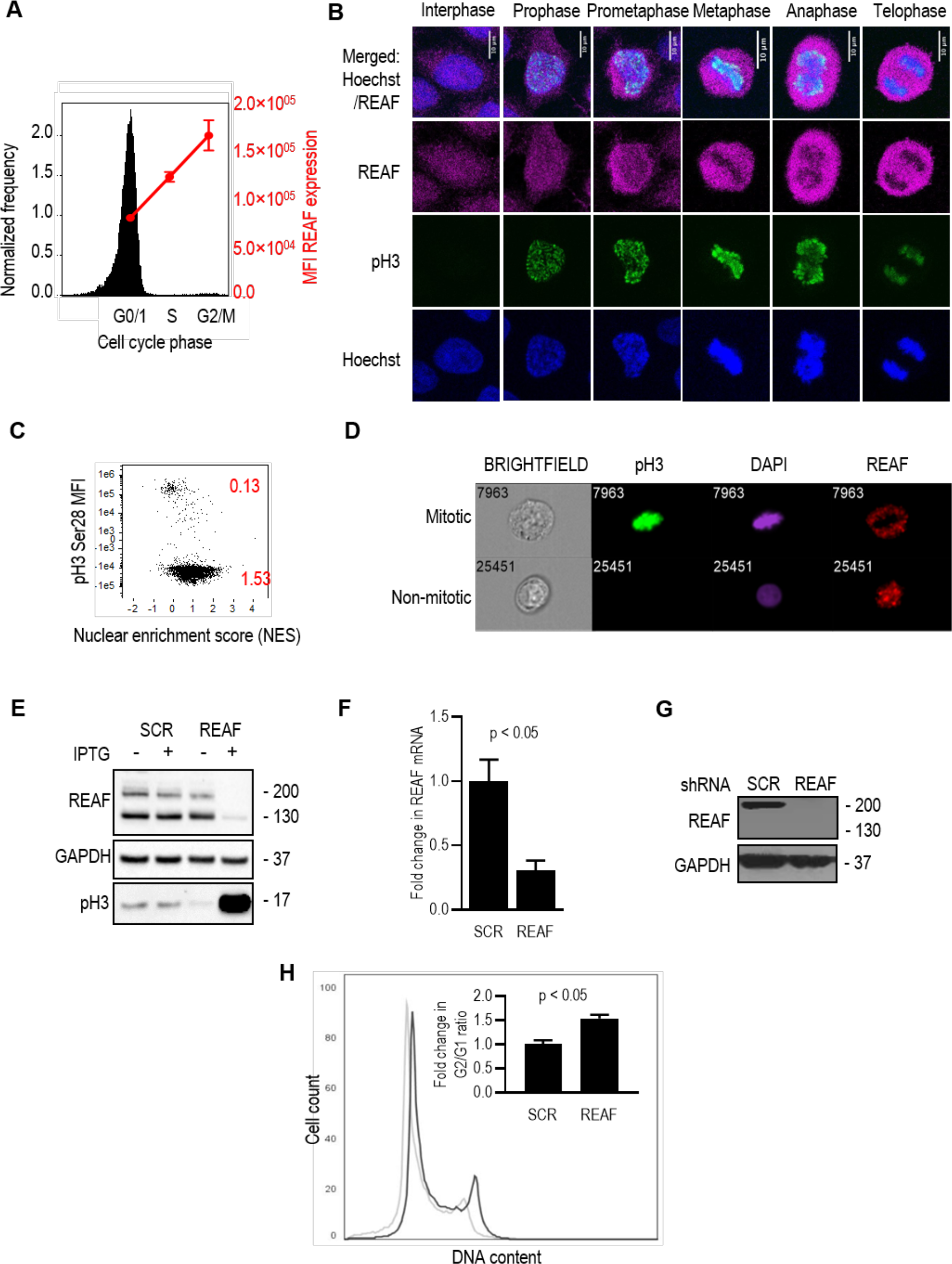
Depletion of REAF results in G2/M accumulation. **(A)** Imaging flow cytometry of cell cycle phase and REAF protein in DAPI stained primary monocytes **(B)** Confocal microscopy of subcellular REAF in HeLa-CD4. Phospho-histone H3 (Ser28) staining and chromatin morphology (Hoechst 33342) were used for cell cycle phase identification. **(C)** Imaging flow cytometry of subcellular REAF in cycling HeLa-CD4. A lower nuclear enrichment score (red) indicates a lower proportion of overall REAF in the nucleus. Phospho-histone H3 (Ser28) staining confirmed mitotic cells had a lower score of 0.13. **(D)** Representative images of subcellular REAF in mitotic and non-mitotic cells. **(E)** REAF protein in THP-1 with IPTG-inducible shRNA targeting REAF (shREAF) or a scrambled control sequence (shSCR). Phospho-histone H3 (Ser10/Thr11) is a mitotic marker and GAPDH is a loading control. (**F)** Fold change in mRNA transcript level in PM1 shREAF normalized to PM1 shSCR measured by qPCR **(G)** REAF protein in PM1 expressing shRNA targeting REAF (shREAF) and PM1 expressing a scrambled control sequence (shSCR). GAPDH is a loading control. **(H)** Flow cytometry of cell cycle phase in PI stained PM1 shREAF (black outline) and PM1 shSCR (grey outline). Plot shown is representative of three biological replicates. Insert shows fold change in G2/G1 ratio in PM1 shREAF normalized to PM1 shSCR. Error bars represent standard deviations of means of three biological replicates.

To determine if the G2/M arrest phenotype, induced by Vpr, could be related to its ability to down modulate REAF, we generated inducible THP-1 and PM1 cell lines that upon induction produce shRNA targeting either REAF or a scrambled control sequence (SCR). After knockdown of REAF in THP-1, there was a clear increase in the expression of the mitotic marker, phosphorylated histone H3 (Ser10/Thr11) (Figure 3E). However, when measured more quantitatively by DNA content analysis in PM1, the potency of the G2/M arrest appeared weak compared to the levels previously described (30, 39). REAF down modulation in PM1 was confirmed by a reduction in mRNA and in protein (Figure 3F and G). Cell cycle phase profiles were determined by flow cytometry (Figure 3H). The increase in the G2/G1 ratio of cells with REAF knocked down, although small (Figure 3H, insert), was comparable to other reports where individual reported targets of Vpr were knocked down (14). In agreement with Greenwood *et al.* 2019, we contend that more than one protein may be required to produce the strong Vpr induced G2/M arrest reported (14, 30, 39).

### REAF is not IFN stimulated or under positive selection

IFNα is central to innate immune responses and is known to induce many HIV-1 restriction factors (43, 44). We used RNA microarray analysis to determine if IFNα upregulated REAF mRNA in MDMs. Figure 4A shows that IFNα induced upregulation of many known antiviral genes, including HIV restriction factors APOBEC3G, MX2, Tetherin and Viperin (45)(46)(43) but with little or no upregulation of REAF mRNA. Nevertheless, antiviral factors are also often upregulated in response to pathogen associated molecular patterns (PAMPs). Poly(I:C) is a double-stranded RNA, used to stimulate molecular pattern recognition pathways associated with viral infection. Figure 4B shows that poly(I:C) induces REAF in THP-1, a macrophage cell line. Lipopolysaccharide (LPS), another PAMP which is TLR4 specific (47), also induces the upregulation of REAF expression in PBMCs (Figure 4C).

**Figure 4:**
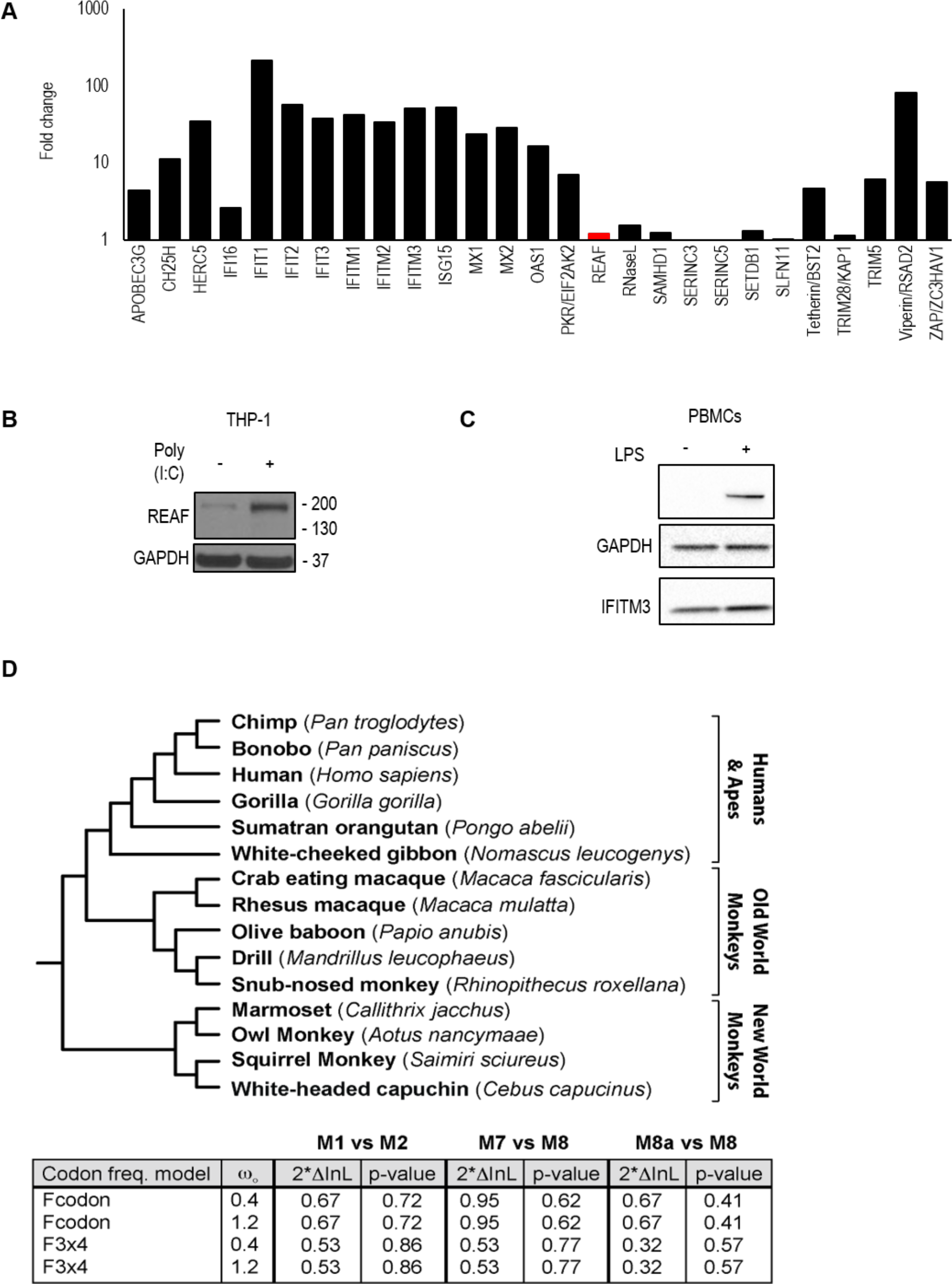
REAF is not IFN stimulated or under positive selection. **(A)** RNA microarray determined change in REAF mRNA compared to other antiviral factors in MDMs treated with IFNα (500IU/ml). **(B)** REAF protein in PMA differentiated THP-1 after poly(I:C) treatment for 48 hours. GAPDH is a loading control. **(C)** REAF protein in PBMCs after LPS treatment. GAPDH is a loading control and IFITM3 is a positive control for LPS induced upregulation. **(D)** REAF DNA sequences from 15 extant primate species (tree length of 0.2 substitutions per site along all branches of the phylogeny) (top) were analyzed using the PAML package for signatures of positive natural selection (bottom). Initial seed values for ω (ω_O_) and different codon frequency models were used in the maximum likelihood simulation. Twice the difference in the natural logs of the likelihoods (2*InL) of the two models were calculated and evaluated using the chi-squared critical value. The p value indicates the confidence with which the null model (M1, M7, M8a) can be rejected in favor of the model of positive selection (M2, M8).

Restriction factors are often under evolutionary positive selection at sites that interact with virus. We compared REAF DNA sequences from 15 extant primate species using PAML package for signatures of positive natural selection. We found no evidence of positive selection of REAF in the primate lineage (Figure 4D).

## Discussion

The deletion of *vpr* in HIV-1 leads to impairment of its replication in both HeLa-CD4 and primary macrophages. A number of experiments presented here point to a role for Vpr in the counter-restriction of the antiviral protein REAF. First, HIV-1 replication is significantly enhanced by knockdown of REAF in either HeLa-CD4 or primary macrophages and this phenotype is more pronounced for viruses lacking *vpr*. Second, REAF is down modulated early after infection in a manner dependent on both the presence of Vpr and, as demonstrated by *vpr* single point mutations, the localization of Vpr to the nuclear envelope and its interaction with a nuclear localised E3 ubiquitin ligase, DCAF1. Third, using VLPs we show that Vpr alone is sufficient to down modulate REAF in MDMs. Finally, by co-immunoprecipitation, we demonstrate that REAF and Vpr physically interact, either directly, or indirectly as part of a complex. Taken together, our results highlight the importance of the relationship between REAF and the HIV-1 accessory protein Vpr.

Others have shown a specific requirement for Vpr in the efficient infection of non-dividing cells and less so in cycling T cells (12, 27, 30). The requirement for Vpr in macrophage infection is substantiated here, reduced viral replication is observed after infection of MDMs with either HIV-1 89.6^Δ*vpr*^ or HIV-1 89.6^F34I^ compared to HIV-1 89.6^WT^. This is the first demonstration of a *vpr*-alleviated impairment of HIV-1 replication in primary macrophages. Recently Yan *et al.* (2019) show that HLTF, a reported target of Vpr, restricts replication of HIV-1 in T cells while Lahouassa *et al.* (2016) also reported a Vpr dependent loss of HLTF at six hours post infection (8, 12). HLTF down modulation occurs concomitantly with REAF as early as 0.5 hours post infection (data not shown). Interestingly, HLTF and REAF were identified in the same screen for proteins that interact with single-stranded DNA (48).

The transient nature and timing of REAF depletion shown here is consistent with its ability to impede the production of reverse transcripts early in infection (20). After an initial down modulation of REAF following infection, REAF depletion is paused, perhaps attributable to the limited quantities of Vpr carried in the virus particle (17). Recently, Greenwood *et al.* carried out a whole cell proteomics screen for factors up or down modulated by Vpr in T cells. They identified almost 2000 proteins affected, underlining the promiscuous activity of Vpr (14). In light of these findings, it is important that attention is directed to those reported Vpr targets that affect replication of HIV-1 in primary cells.

Our model is that Vpr is carried into the cell by HIV-1, in limited, but sufficient quantities to down modulate REAF in the timeframe required for reverse transcription to proceed unhindered. Interestingly, nuclear localisation of Vpr is also required for the down modulation of REAF, perhaps similar to the Vpx mediated depletion of the reverse transcription inhibitor SAMHD1 (for which degradation is initiated in the nucleus) (49). Localisation of Vpr to the nuclear region is a requirement for interaction with REAF and DCAF1 and this results in the degradation of REAF. We propose that REAF is linked to the innate immune response as treatment of cells with poly(I:C) or LPS induces its expression. Furthermore, HIV-1 replication without an intact *vpr*, induces the expression of REAF to high levels in primary macrophages as early as 30 minutes post infection.

A poorly understood event is that Vpr induces cell cycle arrest in the G2/M phase after infection. We report here that the loss of REAF from cycling cells contributes to an accumulation of the population in G2/M. However, the levels of G2/M induction are weak compared to early reports (30, 39). Thus, we contend that Vpr-induced knock down of more than one protein by Vpr may be required for the complete induction of G2/M arrest seen previously. In support of this, G2/M arrest was only weakly induced when Greenwood *et al.* independently knocked down several Vpr targeted proteins such as MCM10, SMN1, CDCA2 and ZNF267 (14).

REAF is unlike the evolving HIV restriction factors such as APOBEC3G, SAMHD1, TRIM5 or BST2/Tetherin and is more similar to SERINC3 and 5 which are not under positive selection (50, 51). REAF has many properties of restriction factors (45, 52). It interacts with HIV-1 reverse transcripts, impeding reverse transcription (20). It is germline encoded, constitutively expressed in cells, regulated by the proteasome system, suppressed by an accessory protein, Vpr, and upregulated by poly(I:C) and LPS. Our results support the current model for Vpr activity which is that it induces the degradation of proteins involved in an unknown restriction of HIV-1. We propose that REAF may be a crucial component a Vpr targeted restriction system that is active against HIV-1.

## Materials and Methods

### Ethics Statement

Leucocyte cones, from which PBMCs were isolated, were obtained from the NHS Blood Transfusion service at St. George’s Hospital, London. Donors were anonymous and thus patient consent was not required. The local ethical approval reference number is 06/Q0603/59.

### Cell Lines

HEK-293T (ATCC), PM1, THP-1, C8166, HeLa-CD4 (all NIBSC AIDS Reagents) and HeLa-CD4 shRNA-REAF (previously described) (18) were maintained at 37°C in 5% CO_2_. Cells were cultured in Dulbecco’s Modified Eagle Medium (DMEM, Thermo Fisher) supplemented with fetal bovine serum (FBS) 5-10% and appropriate antibiotics (all Thermo Fisher). HeLa-CD4-shRNA-REAF were selected for resistance to puromycin in media supplemented with 10μg/ml puromycin (Invitrogen).

The isopropyl β-D-1-thiogalactopyranoside (IPTG)-inducible vector pLKO-IPTG-3xLacO (Sigma) was used to express short hairpin RNAs (shRNAs) targeted against REAF (Mission TRCN0000141116, Sigma). Additionally, a non-target (scramble) control was prepared. Viral particles for cell line transductions were prepared by co-transfecting HEK-293T cells with pLKO-IPTG-3xLacO, the Gag/Pol packaging vector pLP1, a Rev expression vector pLP2, and the vesicular stomatitis virus G protein (VSV-G) expression vector pVPack-VSV-G (Stratagene). After 72 hours, virus was clarified by low-speed centrifugation and passed through a 0.45-m-pore-size filter. THP-1 and PM1 cells were transduced by culturing viral particles in the presence of 8g/ml Polybrene for 72 hours, after which resistant colonies were selected and maintained with 2μg/ml puromycin. Culturing cells in the presence of 1mM IPTG for 72 hours induced expression of shRNAs.

### Transfections and Virus/VLP Production

The infectious molecular clone for HIV-1 89.6 was obtained from the Centre for AIDS Research (NIBSC, UK). Infectious full-length and chimeric HIV clones were prepared by linear polyethylenimine 25K (Polysciences), Lipofectamine 2000 (Invitrogen) or Lipofectamine 3000 (Invitrogen) transfection of HEK-293T. Virus-like particles (VLPs) were produced by linear polyethylenimine 25K (Polysciences) transfection of HEK-293T. The VLP packaging vector was a gift from N. Landau and production is described in reference (27).

The plasmid construct HIV-1 89.6^Δ*vpr*^ was generated from the HIV-1 89.6 molecular clone, using overlap extension PCR *(44)*. Clones were confirmed by plasmid sequencing (Source BioScience). Primer sequences are available upon request. HIV-1 p89.6 *vpr* mutants F34I and Q65R were made by site directed mutagenesis (Agilent) of the p89.6 plasmid. HEK-293T were plated at 2×10^4^/cm^2^ in 10cm dishes (for virus and VLP production) 48 hours prior to transfection. For virus/VLP production, supernatant was harvested 72 hours post transfection and cleared of cell debris by centrifugation at 500 × *g* for 5 minutes. All viruses were amplified by C8166 for 48 hours prior to harvest.

### Titration of Replication Competent Virus

HeLa-CD4 were seeded at 1.5×10^4^ cells/well in 48-well plates to form an adherent monolayer of cells. Cell monolayers were challenged with serial 1/5 dilutions of virus and titre was assessed after 48 hours by *in situ* intracellular staining of HIV-1 p24 to identify individual foci of viral replication (FFU), as described previously (53). For infection time course experiments, 400-500μl of 1×10^5^ FFU/ml (HeLa-CD4) or 3×10^3^ FFU/ml (MDMs) virus was added per well to cells cultured in 6-well trays for 24 hours (HeLa-CD4) or 7 days (for MDMs). For Figure 2F and 2H, cells were challenged with 50ng p24 in 6-well plates with 2×10^6^ MDMs per well. For Figure 2F, supernatants were harvested on days 0, 2, 8, 21 and 28 post challenge and p24 concentration analysed by ELISA.

### p24 ELISA

ELISA plates were pre-coated with 5μg/ml sheep anti-HIV-1 p24 antibody (Aalto Bio Reagents) at 4°C overnight. Viral supernatants were treated with 1% Empigen^®^ BB for 30 minutes at 56°C, then plated at 1:10 dilution in tris-bufered saline (TBS) on pre-coated plates and incubated for 3 hours at room temperature. Alkaline phosphatase-conjugated mouse anti-HIV-1 p24 monoclonal antibody (Aalto Bio Reagents) in TBS 20% sheep serum, 0.05% v/v Tween-20 was then added and incubated for 1 hour at room temperature. After 4 washes with PBS 0.01% v/v Tween-20 and 2 washes with ELISA Light washing buffer (ThermoFisher), CSPD substrate with Sapphire II enhancer (ThermoFisher) was added and incubated for 30 minutes at room temperature before chemiluminescence detection using a a plate reader.

### cDNA Synthesis and qPCR

Total RNA was extracted from PM1 cells using the ReliaPrep RNA Kit (Promega). One-step reverse transcription qPCR (Quantbio) using TaqMan probes detected amplified transcripts. Data acquired by an Agilent Mx3000 was analyzed with MxPro software.

### Gene Expression RNA microarray

Prior to microarray analysis, RNA from MDMs was prepared using the Illumina™ TotalPrep™ RNA Amplification Kit (Ambion), according to manufacturer’s instructions. The probes were hybridized on an Illumina™ HT12v3 bead array following the manufacturer’s standard hybridization and scanning protocols. Raw measurements were processed by GenomeStudio software (Illumina), and quantile normalized. Microarray data are publicly available in the Gene Expression Omnibus (GEO) database with accession number GSE54455.

### IFN, Poly(I:C) and LPS Treatment

MDMs were treated with IFN (500IU/ml) for 24 hours before harvest for RNA extraction. Recombinant IFNα was purchased from Sigma (Interferon-αA/D human Cat. No. I4401-100KU) and is a combination of human subtypes 1 and 2. THP-1 were treated with poly(I:C) (25μg/ml, HMW/LyoVec™, Invitrogen) for 48 hours before analysis by Western blotting. Prior to poly(I:C) treatment, THP-1 were treated with phorbol 12-myristate 13-acetate (PMA, 62ng/ml) for 3 days and then PMA-free DMEM for 2 days to allow differentiation and recovery. PBMCs isolated from healthy blood donors were treated with lipopolysaccharide (LPS, 10 ng/ml) for 24 hours before analysis by Western blotting.

### Western blotting

Cells were harvested and lysed in 30-50μl of radioimmunoprecipitation assay (RIPA) buffer supplemented with NaF (5μM), Na_2_VO_3_ (5μM), β-glycerophosphate (5μM) and 1x Protease Inhibitor Cocktail (Cytoskeleton). The protein concentration of each sample was determined using BCA Protein Assay Kit (Pierce). 12.5-70μg of total protein was separated by SDS-PAGE (4-12% Bis-Tris Gels, Invitrogen), at 120V for 1 hour 45 minutes in MOPS SDS Running Buffer (Invitrogen). Separated proteins were transferred onto nitrocellulose membrane (0.45μm pore size, GE Healthcare) at 45V for 2 hours, in ice-cold 20% (v/v) Methanol NuPAGE™ Transfer Buffer (ThermoFisher). After transfer, membranes were stained for total protein using Ponceau S staining solution (0.1% (w/v) Ponceau in 5% (v/v) acetic acid), washed 3 times for 5 minutes on an orbital shaker in dH_2_O and imaged using ChemiDoc Gel Imaging System. Membranes were blocked for 1 hour at room temperature in 5% (w/v) non-fat milk powder in TBS-T buffer. Specific proteins were detected with primary antibodies by incubation with membranes overnight at 4°C and with secondary antibodies for 1 hour at room temperature. All antibodies were diluted in blocking buffer. Proteins were visualized using ECL Prime Western Blotting Detection Reagent (GE Healthcare) and imaged using either ChemiDoc Gel Imaging System (Bio-Rad) or exposed to CL-XPosure films (ThermoScientific) and developed. In all places where quantitative comparisons are made, such as in Figures 1D and E, blots are derived from the same blot or blots processed together.

### Antibodies

Primary rabbit polyclonal antibody to REAF (RbpAb-RPRD2) has been previously described (20). For imaging flow cytometry and confocal microscopy, RbpAb-RPRD2 was detected using goat anti-rabbit IgG conjugated with Alexa Fluor 647 (Invitrogen). FITC-labelled anti-phospho-histone H3 (Ser28) Alexa 488 was used (BD Bioscience) for imaging flow cytometry and confocal microscopy. MsmAb-GFP (both Abcam) was detected by anti-mouse IgG antibody conjugated to HRP (GE Healthcare) for Western blotting. Also for Western blotting, RbpAb-RPRD2, RbmAb-IFITM3 (EPR5242, Insight Biotechnology), RbpAb-GAPDH, and RbmAb-phospho-histone H3 (Ser10/Thr11) were detected with donkey anti-rabbit IgG conjugated to HRP (GE Healthcare).

### Immunoprecipitation

HEK-293T, transfected with either Vpr-GFP expression plasmid or GFP control expression vector, were lysed 72hrs post transfection in RIPA buffer supplemented with NaF (5μM), Na_2_VO_3_ (5μM), β-glycerophosphate (5μM) and 1x Protease Inhibitor Cocktail (Cytoskeleton). Total protein concentration was determined using BCA Protein Assay Kit (Pierce). GFP-TRAP^®^ magnetic agarose beads were equilibrated in ice cold dilution buffer (10 mM Tris/Cl pH 7.5; 150 mM NaCl; 0.5 mM EDTA) according to manufacturer’s instructions (Chromotek). Cell lysates containing 100μg of total protein were incubated with 10μl of equilibrated beads for 2 hours at 4°C with gentle agitation. Beads were washed three times with PBST buffer before analysis of immunoprecipitated protein by Western blotting.

### Magnetic Separation of Primary Human Lymphocytes

Peripheral blood mononuclear cells (PBMCs) were isolated from leukocyte cones (NHS Blood Transfusion service, St. George’s Hospital, London) by density gradient centrifugation with Lymphoprep™ density gradient medium (STEMCELL™ Technologies). Peripheral monocytes were isolated from PBMCs, using the human CD14^+^ magnetic beads (Miltenyi Biotech) according to manufacturer’s instructions. CD4^+^ T cells were isolated from the flow-through, using the human CD4^+^ T cell isolation kit (Miltenyi Biotech). CD14^+^ monocytes, and CD4^+^ T cells were either differentiated, or fixed directly after isolation for intracellular staining. To obtain M1 and M2 macrophages (M1/M2 MDMs), monocytes were treated with either granulocyte-macrophage colony stimulating factor (GM-CSF, 100ng/ml, Peprotech) or macrophage colony stimulating factor (M-CSF, 100ng/ml) for 7 days, with medium replenished on day 4. To obtain dendritic cells (DC), monocytes were treated with GM-CSF (50ng/ml) and IL-4 (50ng/ml) for 7 days, with medium replenished on day 4. Activated CD4^+^ T cells were obtained by stimulating freshly isolated CD4^+^ T cells at 1×10^6^/ml with T cell activator CD3/CD28 Dynabeads (ThermoFisher), at a bead-cell-ratio of 1, for 7 days. Magnetic beads were removed prior to intracellular staining and imaging flow cytometry.

### Immunofluorescence

HeLa-CD4 were plated at 2×10^4^/cm^2^ in 8-well chamber slides for confocal microscopy. Cells were washed with PBS, fixed in 2% paraformaldehyde/PBS for 10 minutes at room temperature. Fixed cells were permeabilized in 0.2% Triton™-X100/PBS for 20 minutes at room temperature and incubated with primary antibodies in PBS 0.1% Triton-X100 2% BSA overnight at 4°C. After 3 washes in PBS, cells were labelled with secondary antibodies in the same buffer for 1 hour at room temperature, and washed 3 times with PBS. Nuclei were counterstained with Hoechst 33342 (2μM, ThermoFisher) for 5 minutes at room temperature. Labelled cells were mounted with ProLong™ Diamond Antifade Mountant (ThermoFisher) and analyzed on a laser scanning confocal microscope LSM 710 (Carl Zeiss). Images were acquired with ZEN software and analyzed with ImageJ.

### Imaging Flow Cytometry

Cells were fixed in FIX&PERM^®^ Solution A (Nordic MUbio) for 30 minutes, and permeabilized with 0.2% Triton™-X 100/PBS. MDMs were blocked with human serum (1%). The staining buffer used was: 0.1% Triton™-X 100 0.5% FBS. Nuclei were counterstained with DAPI (1μg/ml) for two hours. Imaging flow cytometry was performed using the Amnis ImageStream^®x^ Mark II Flow Cytometer (Merck) and INSPIRE^®^ software (Amnis). A minimum of 5000 events were collected for each sample. IDEAS^®^ software (Amnis) was used for analysis and to determine the ‘nuclear enrichment score’ (NES). The NES is a comparison of the intensity of REAF fluorescence inside the nucleus (defined using the exclusively nuclear stain DAPI) to the total fluorescence intensity of REAF in the entire cell (defined using brightfield images). A lower nuclear enrichment score indicates a lower proportion of overall REAF is located within the nucleus.

### Statistics

Statistical significance in all experiments was calculated by Student’s t-test (two tailed) or ANOVA (indicated). Data are represented as mean ± standard deviation (error bars). GraphPad Prism and Excel were used for calculation and illustration of graphs.

### Cell Cycle Analysis

Cell cycle phase distribution was determined by analysis of DNA content via either flow cytometry (BD FACS Canto™ II) or imaging flow cytometry. Cells were fixed in FIX&PERM^®^ Solution A (Nordic MUbio) and stained with DAPI (1μg/ml) before analysis by imaging flow cytometry. Cell lysates were assessed by Western blotting using the anti-phospho-histone H3 (Ser10/Thr11) antibody as an additional mitotic marker. Chromatin morphology and anti-phospho-histone H3 (Ser28) were used to determine the cells in indicated phases of the cell cycle and mitosis in confocal microscopy experiments. Cell cycle status of PM1 cells was determined via propidium iodide (PI) staining using FxCycle PI/RNse solution (ThermoFisher). Stained cells were analyzed on an NxT flow cytometer (ThermoFisher).

### Evolutionary Analysis

To ascertain the evolutionary trajectory of REAF, we analyzed DNA sequence alignments of REAF from 15 species of extant primates using codeml (as implemented by PAML 4.2) (54). The evolution of REAF was compared to several NSsites models of selection, M1, M7 and M8a (neutral models with site classes of dN/dS <1 or ≤1) and M2, M8 (positive selection models allowing an additional site class with dN/dS >1). Two models of codon frequencies (F61 and F3×4) and two different seed values for dN/dS (ω) were used in the maximum likelihood simulations. Likelihood ratio tests were performed to evaluate which model of evolution the data fit significantly better. The p-value indicates the confidence with which the null model (M1, M7, M8a) can be rejected in favor of the model of positive selection (M2, M8). The alignment of REAF was analyzed by GARD to confirm the lack recombination during REAF evolution (55). Neither positively selected sites nor signatures of episodic diversifying selection were detected within REAF by additional evolutionary analysis by REL and FEL or MEME (56).

### Data Availability

All RNA microarray data is available in the gene expression omnibus (GEO) database with accession number GSE54455.

## Acknowledgements

This work was supported partly by an MRC Senior Non-Clinical Fellowship awarded to AM (G117/547) and PhD studentships awarded by QMUL Life Sciences Institute (LSI) (CEJ) and The Rosetrees Trust (JMG and CEJ, M665 and M275). RDS was supported by the Wellcome Trust-University of Edinburgh Institutional Strategic Support Fund. The monoclonal antibodies to HIV-1 p24 (EVA365 and 366) were provided by the EU Programme EVA Centre for AIDS Reagents, NIBSC, UK (AVIP Contract Number LSHP-CT-2004-503487). The Wellcome Trust (101604/Z/13/Z) funded the purchase of Amnis ImageStream™ imaging flow cytometer. We wish to thank N. Landau for the kind gift of VPL constructs.

